# Amygdala metabolic activity on [^18^F]FDG PET is associated with survival, response to immune checkpoint inhibition, and tumor immune signaling in non-small cell lung cancer

**DOI:** 10.64898/2026.09.10.750685

**Authors:** Barbara Katharina Geist, Chrysoula Vraka, Clara T.S. Mertel, Lucian Beer, Armin Frille, Laurin Herbsthofer, Nadine Hermann, Swen Hesse, Lukas Kenner, Daria Kifjak, Kilian Kluge, Oana Cristina Kulterer, Constantin Lapa, Alessandro Lieblich, Thomas Selim Nakuz, Felicitas Oberndorfer, Renate Kain, Helmut Prosch, Osama Sabri, Bernhard Sattler, Thomas Schweiger, Clemens Aigner, Clemens Spielvogel, Dato Tsomaia, Karolina Trachtova, Martina Tomberger, Hubert Wirtz, Josef Yu, Stefan Grünert, Marcus Hacker

## Abstract

Amygdala metabolic activity (AmygAc) measured by [^18^F]FDG PET has been associated with chronic psychological stress and adverse clinical outcomes, but its relationship with antitumor immunity and treatment response remains incompletely understood. We investigated AmygAc as a host-derived imaging biomarker in non-small cell lung cancer (NSCLC) and examined its association with survival, response to immune checkpoint inhibition (ICI), and the tumor immune microenvironment.

**Methods:** In this multicenter retrospective study, baseline [^18^F]FDG PET scans from 256 treatment-naïve patients with NSCLC were analyzed to quantify AmygAc. Overall survival was assessed in the full cohort, and treatment response was evaluated in 79 patients who subsequently received neoadjuvant ICI therapy. AmygAc was integrated with clinical and tumor-derived imaging parameters. To investigate its biological correlates, tumor tissue from a selected subcohort was analyzed using multiplex immunofluorescence and spatial transcriptomics.

**Results:** Higher AmygAc was associated with poorer survival and remained independently associated with outcome after multivariable adjustment for established clinical and imaging parameters. Among patients receiving neoadjuvant ICI therapy, AmygAc was lower in responders than in non-responders (0.76 vs. 0.81, *P* = 0.003) and discriminated treatment response with an area under the receiver operating characteristic curve of 0.80. Molecular profiling demonstrated attenuated interferon-γ signaling in tumors from patients with higher AmygAc (normalized enrichment score = −1.65; false discovery rate = 0.015). Multiplex immunofluorescence further showed a non-significant trend toward reduced cytotoxic T-cell infiltration together with significantly increased spatial proximity of cytotoxic T cells to PD-L1-expressing cells, without corresponding differences in overall PD-L1 expression.

**Conclusions:** AmygAc derived from routine [^18^F]FDG PET is a host-derived imaging biomarker associated with survival, ICI response, and distinct features of the tumor immune microenvironment in NSCLC. These findings link neural metabolic activity with antitumor immune regulation and support prospective evaluation of AmygAc as a complementary biomarker for risk stratification and immunotherapy response.

## Introduction

Functional imaging using Positron Emission Tomography (PET) with the glucose analogue 2-deoxy-2[^18^F]fluoro-D-glucose ([^18^F]FDG) enables the detection of metabolically active cells at ectopic sites throughout the entire body. The [^18^F]FDG-PET derived imaging biomarker total lesion glycolysis (TLG) has demonstrated prognostic value in oncology [1,2] and has become a cornerstone for stratification of cancer patients. Owing to its non-specific uptake beyond malignant cells, [^18^F]FDG PET is also widely used for imaging infection, inflammation, and a broad spectrum of neurological disorders. Neuroscience pioneered metabolic network and functional connectivity analyses of the brain using specialized fMRI and functional PET protocols [3]. More recently, advances in long-axial field-of-view (LAFOV) PET, and artificial intelligence (AI)-enabled computational methods have enabled a paradigm shift in whole-body imaging toward comprehensive assessment of systemic metabolic interactions across organs [4].

In this context, the broad biodistribution of [^18^F]FDG reflecting glucose metabolism throughout the body is a unique strength, rather than a limitation. This systems-level perspective extends the clinical utility of [^18^F]FDG PET beyond established total lesion glycolysis (TLG), and beyond conventional analyses of brain connectivity [3].

Altered [^18^F]FDG uptake in the bone marrow and spleen reflect systemic inflammatory activity in cancer and has been associated with treatment-related toxicity, therapeutic response, and clinical outcome [5–7]. A global analysis of whole body FDG-PET scans revealed parameters like density and structure of organ connectomes correlate with outcome in cancer patients, however these tools remain research oriented. However, currently no imaging biomarkers outside the lesion has entered clinical use, due to a variety of technical challenges, including variability and harmonisation across multiple sites.

Within the central nervous system, Yang and colleagues demonstrated widespread alterations in brain glucose metabolism across different cancer types and disease stages, with the most pronounced metabolic changes observed in patients with lung cancer[8]. Among brain-derived imaging biomarkers, elevated amygdala metabolic activity (AmygAc) has emerged as a robust marker of chronic psychological stress, depression, and anxiety. Increased AmygAc has been shown to predict stress-related cardiovascular events, survival in patients with head and neck cancer, and postmenopausal osteoporosis [9–11]. More recently, AmygAc has also been associated with multi-organ metabolic profiles and response to neoadjuvant immunochemotherapy in patients with non-small cell lung cancer (NSCLC)[12]. Although previous studies have established the prognostic value of AmygAc, the molecular mechanisms linking elevated stress-related neural activity to metabolic reprogramming, remodeling of the tumor immune microenvironment (TIME), and adverse clinical outcomes remain poorly understood. Consequently, the biological basis underlying its prognostic and predictive significance has yet to be defined.

Here, we conducted a multi-center retrospective study of patients with NSCLC to evaluate the readily accessible AmygAc in routine FDG-PET scans as a host-derived imaging biomarker and investigate its association with the TIME, outcomes and influence on the response to immune checkpoint inhibitor (ICI) therapy. By integrating large field of view (LAFOV) metabolic imaging, molecular profiling, and clinical outcome data, we reveal a robust relationship between elevated AmygAc and adverse outcome in cancer patients, which is related to tumor metabolism, immune activation, and response to neoadjuvant immune checkpoint inhibition.

## Methods

### Study design and patient cohort

The retrospective cohort from the Vienna General Hospital (AKH) was approved by the Austrian authorities and ethics committee at the Medical University of Vienna (no. 1649/2016), including the usage of pseudonymized PET/CT and patient data in conjunction with surgically removed cancer tissue for phenotyping and genetic analyses. From a pool of 1148 lung cancer patients undergoing a PET/CT between 2014 and 2018, 174 therapy naïve subjects with whole body scan (including the entire head) were included, of which we could access 20 samples for multiplexing immunofluorescence (MP-IF) imaging, genetic and transcriptomic analyses and from another 38, follow-up CT scans after surgery, chemo or radio therapy were available, for details see Figure 1. Another retrospective cohort was available from Klinik Floridsdorf in Vienna, from which 19 patients were included, receiving whole-body PET/CT scans between 2022 and 2024 (ethics approval no. 1521/2015). All of them received neoadjuvant immune checkpoint inhibitor (ICI) therapy after the baseline scan and they were followed for at least one year. Identically, from University of Augsburg, Germany, five patients were included with whole-body PET/CT scans and neoadjuvant immune checkpoint inhibitor (ICI), being followed for at least one year. And from Leipzig

**Figure 1:**
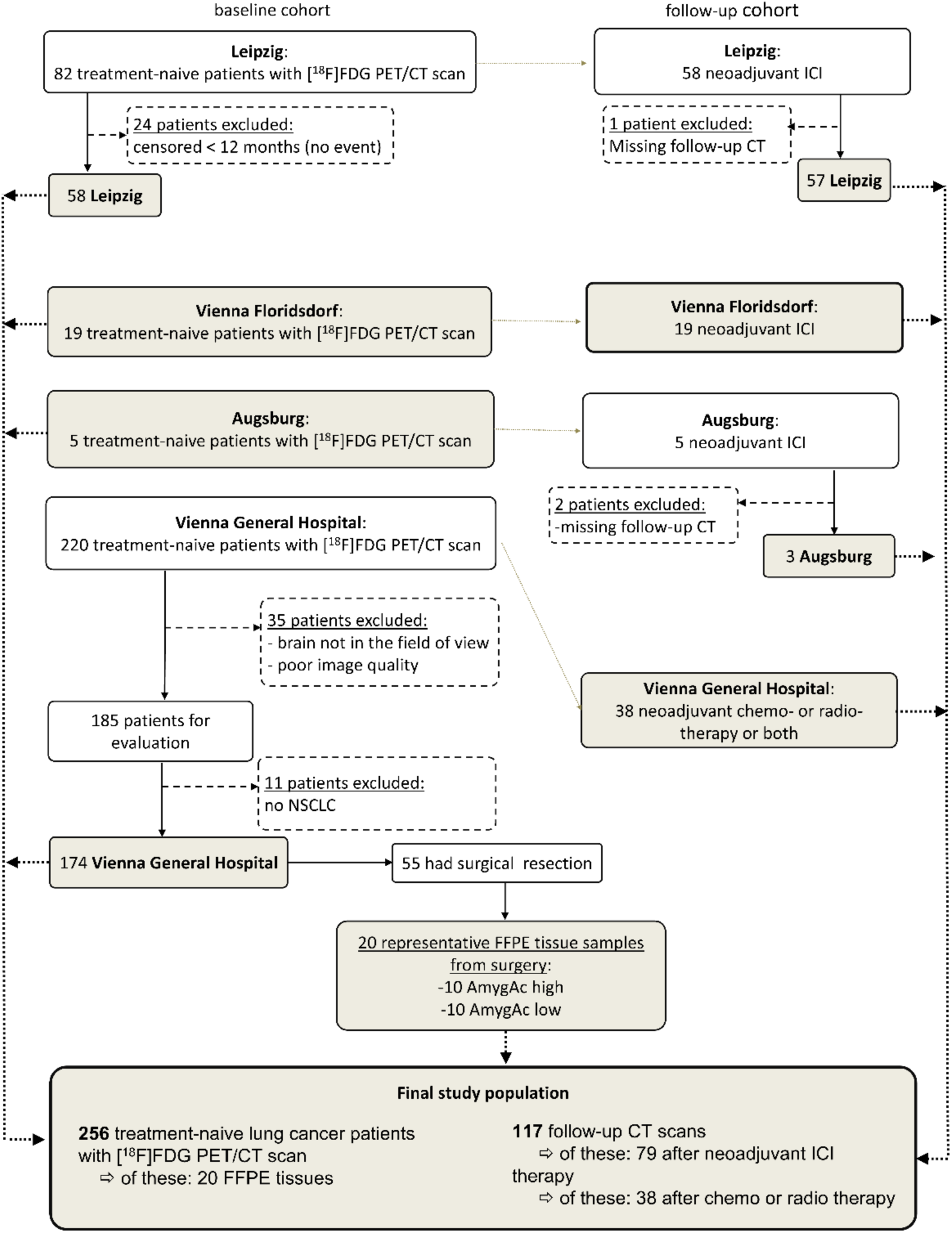
Study flow diagram and cohort composition of the study.

University Medical Center, Germany, another 58 patients, from which biopsies were used to analyze PD-L1 levels in tumor and immune cells.

From each patient, blood values including c-reactive protein (CRP) and leucocytes at the time of the baseline scan were collected as well as demographics, tumor stage, follow-up data and survival status, see table 1.

**Table 1:** Baseline demographic, clinical and PET-based parameters stratified for patients with and without 1 year survival. Values are presented as mean (standard deviation) for the full cohort and the follow-up sub-cohort (separated with backslash). AC, Adenocarcinoma; SSC, Squamous Cell Carcinoma; BMI, Body Mass Index; CRP, C-Reactive Protein; ICI, Immune Checkpoint Inhibitor; NSCLC, Non-Small-Cell Lung Cancer;

|  |  |  |
| --- | --- | --- |
|  | Full cohort | Follow-up cohort |

|  | Total cohort | survived 1st year | Death within 1st year | P-value | Total cohort | survived 1st year | Death within 1st year | P-value |
| --- | --- | --- | --- | --- | --- | --- | --- | --- |
| Clinical and demographic parameters |  |  |  |  |  |  |  |  |
| Subjects (% male) | 256 (50 %) | 183 (49 %) | 73 (52 %) |  | 117 (43 %) | 65 (60 %) | 52 (21 %) |  |
| Age [years] | 65 (10) | 65 (10) | 65 (12) | 0.99 | 62 (9) | 61 (9) | 62 (9) | 0.4 |
| BMI [kg/m<sup>2</sup>] | 26 (5) | 26 (5) | 25 (5) | 0.5 | 25 (4) | 25 (3) | 25 (4) | 0.9 |
| Leucocytes [10<sup>9</sup>/l] | 9.2 (2.8) | 9.2 (2.7) | 9.1 (3.0) | 0.9 | 10.2 (1.9) | 10.8 (2.2) | 9.9 (1.9) | 0.5 |
| CRP [mg/dl] | 4.4 (8.4) | 4.6(9.3) | 3.8 (5.1) | 0.6 | 11.4 (15.9) | 11.8 (22.5) | 11.3 (13.0) | 0.9 |
| Tumor related parameters |  |  |  |  |  |  |  |  |
| Histology: |  |  |  |  |  |  |  |  |
| AC | 113 | 80 | 33 | 0.6 | 15 | 10 | 5 | 0.4 |
| SCC | 64 | 46 | 18 |  | 10 | 6 | 4 |  |
| Non-defined NSCLC | 79 | 57 | 22 |  | 231 | 49 | 43 |  |
| Total lesion glycolysis TLG | 531 (242) | 340 (588) | 905 (1072) | < 0.001 | 512 (762) | 256 (480) | 698 (875) | 0.03 |
| Stages: |  |  |  | < 0.001 |  |  |  | 0.003 |
| Stage I | 32 | 29 | 3 |  | 0 |  |  |  |
| Stage II | 21 | 16 | 5 |  | 5 | 5 | 2 |  |
| Stage III | 73 | 61 | 12 |  | 23 | 21 | 20 |  |
| Stage IV | 120 | 68 | 52 |  | 44 | 24 | 22 |  |
| Radiotherapy |  |  |  |  | 1 |  |  |  |
| Chemotherapy |  |  |  |  | 38 | 35 | 3 | 0.01 |
| Surgery |  |  |  |  | 24 | 20 | 4 | 0.1 |
| ICI therapy |  |  |  |  | 79 | 56 | 23 | 0.01 |
| Pembrolizumab |  |  |  |  | 55 | 39 | 16 | 0.8 |
| Nivolumab |  |  |  |  | 12 | 10 | 2 |  |
| Durvalumab |  |  |  |  | 3 | 2 | 1 |  |
| Atezolizumab |  |  |  |  | 4 | 2 | 2 |  |
| PET-derived bio markers (mean TBR) |  |  |  |  |  |  |  |  |
| amygdala | 0.84<br>(0.11) | 0.82 (0.11) | 0.87 (0.12) | 0.002 | 0.83 (0.11) | 0.84 (0.11) | 0.84 (0.11) | 0.9 |
| Bone marrow | 1.4 (1.1) | 1.4 (1.1) | 1.6 (1.2) | 0.3 | 1.7 (1.0) | 1.7 (0.9) | 1.6 (0.9) | 0.6 |

### PET/CT image acquisition and imaging outcomes

All patients had fasted a minimum of 6 hours, blood glucose value was measured prior to tracer injection of 250-300 MBq [^18^F]FDG and they rested in a calm environment recumbent afterwards. 60 minutes after injection, images were obtained from the skull to the upper femur at the Vienna General Hospital (AKH) as well as at Klinik Floridsdorf as well as in Augsburg on a Siemens Biograph TruePoint 64 PET/CT system (Siemens Healthineers, Erlangen, Germany). In Augsburg, two patients were additionally scanned on a GE HealthCare Discovery MI. PET scans were acquired in 3-4 bed positions with 3 minutes per bed position followed by a CT scan at 120 kVp and 100 mAs with intravenous contrast, which was also used for attenuation correction. PET images on Siemens scanners were reconstructed iteratively using the ordered-subset expectation maximization (OSEM) algorithm four iterations per 21 subsets, a 5mm slice thickness and a 168 x 168 matrix. PET images on the GE scanner were reconstructed with a matrix size of 256 × 256 × 359 using a Bayesian penalized likelihood reconstruction algorithm (Q.Clear) with a beta value of 600, PSF + TOF and all corrections applied.

Image analysis was performed by altogether four nuclear medicine physicians on a dedicated workstation using the Hybrid 3D software (version 4, Hermes Medical Solutions, Stockholm, Sweden). Amygdala and brain background activity were acquired as previously described [13]: for the amygdala activity, a cubic volume of interest (VOI) was placed in the left amygdala, mean and maximum tracer concentrations were measured as body weight corrected standardized uptake value (SUV). Brain background activity was determined from the temporal lobe SUV_mean_. The left amygdala activity expressed in SUV_mean_ and SUV_max_ were normalized to the temporal lobe background SUV_mean_, thus the so-called signal-to-noise-ratios SNR_mean_ and SNR_max_ served as primary and secondary markers for chronic stress, respectively. For the results of the secondary marker see supplement.

Tumor volumes were delineated semi-automatically using liver segment VIII as reference region. Automatically generated non-tumor organs with high [^18^F]FDG uptake (e.g. bladder or brain) were manually removed from the tumor mass. Mean tumor volume (MTV) and tumor SUV_mean_ were measured, and their product, the total lesion glycolysis (TLG) was further used as a marker of tumor metabolic activity.

[^18^F]FDG uptake in the bone marrow was measured as maximum SUV between lumbar vertebrae L1 and L5. Blood pool activity was obtained through a representative intravascular VOI in the ascending aorta, according to an established method [14]. Similarly, bone marrow was obtained through representative VOIs, adapted from a previously described method[15]. Bone marrow SNR_max_, normalized to the blood pool SUV_mean_ (thus bone marrow SNR_max_), were investigated and served as imaging marker of inflammatory activation. Representative images are depicted in Figure 2.

**Figure 2:**
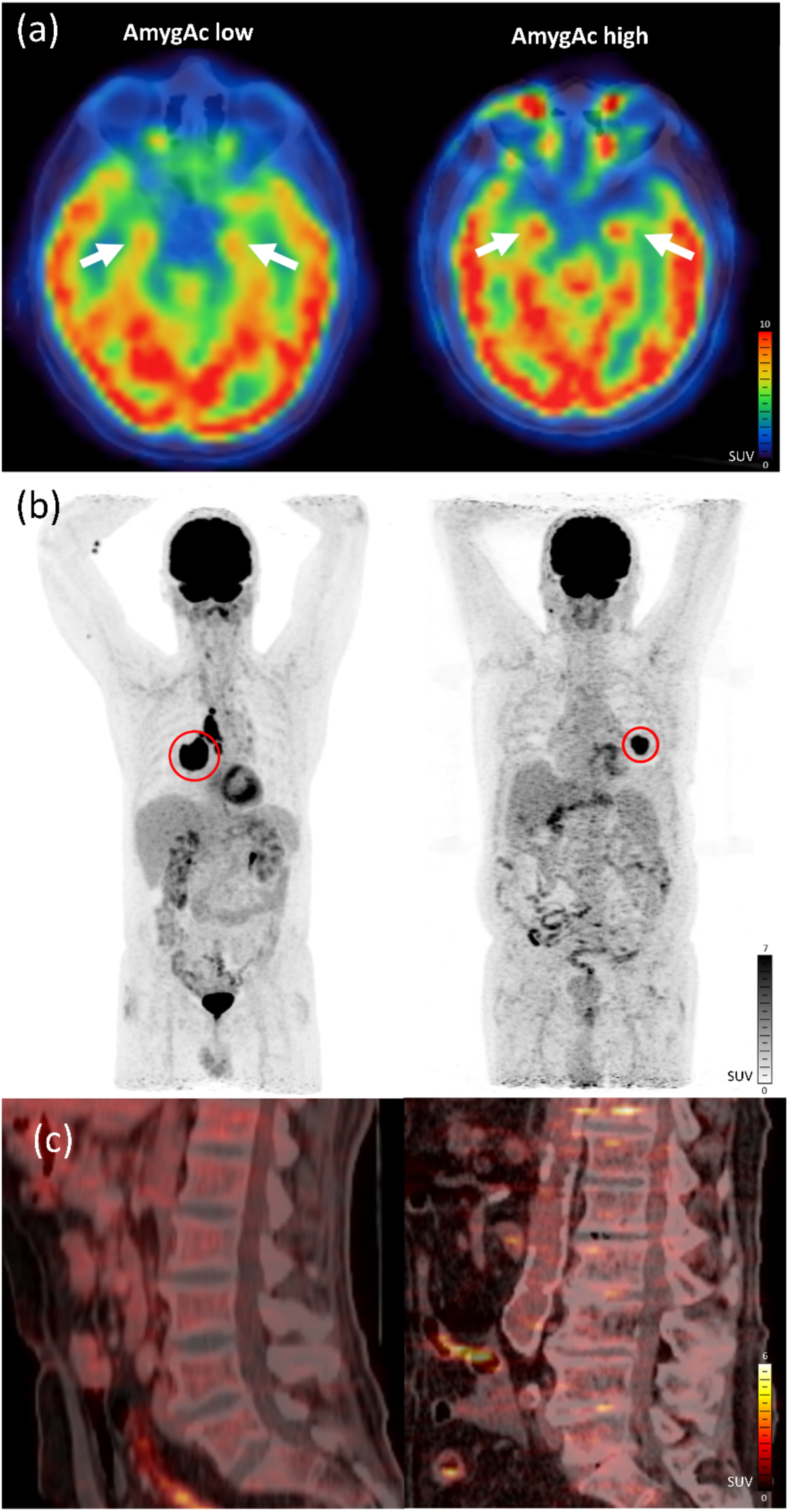
Representative examples of whole body [^18^F]FDG PET scans in patients with low (left panel) and high (right panel) amygdala activity; (a) axial slices through the brain with amygdala region indicated with white arrows; (b) whole body maximum-intensity-projections with primary tumors marked in the red circles; (c) sagittal PET/CT sliced demonstrating increased [^18^F]FDG bone marrow uptake in the lumbar region in the patient with increased amygdala activation. [^18^F]FDG PET activity is quantified with the standardized uptake value (SUV).

In the follow-up cohort, therapy response was classified on the volumetric response assessment on basis of the RECIST guidelines [16] by calculating the tumor volume from the follow-up CTs. The cohort was then dichotomized into responders, i.e. patients with complete or partial response (at least 30 % reduction of tumor volume) and non-responders (all others).

### Statistics Clinical Data

Statistical evaluations were performed using SPSS (version 27.0.1.0), the survival-ROC analyses with a Kaplan-Meier estimator [17] were performed with R (version 4.1.0). The mediation analysis was performed with the PROCESS macro for SPSS from A. F. Hayes [18], using model 6 with 10000 bootstrap samples. The mediation analysis was based on regressions to estimate direct and indirect effects and produced confidence intervals. A P-value of less than 0.05 was classified as statistically significant. All variables were tested for normal distribution with a Shapiro-Wilk test and by inspection of the histograms.

### Patient tumor tissue for molecular analyses

The 20 samples were matched for gender and included balanced numbers of histological tumor types for all respectively analysis. The recorded classifications were confirmed from a pathologist and classified for the histological phenotypes squamous cell carcinoma (SCC, n = 10), adenocarcinoma (AC, n = 8) as well as large cell carcinoma (LCC) and adenosquamous carcinoma (ADSQ; both n = 1). Tumor area (characterized by a high density of tumor cells), TIME (characterized by disordered tissue with immune cell infiltration) and healthy tissue (parenchyma albeit highly infiltrated by immune cells) were delineated and ambiguous samples were clarified in consultation with a pathologist.

### Genetics

The same 20 tissue samples (ten per stress group), which were used for multiplexing IF and spatial transcriptomics, were taken for full genomic sequencing and evaluation of genes most frequently altered in lung cancer (e.g. EGFR, KRAS, P53, BRAF, ROS1, ALK etc.)[19,20]. For full sequencing heterogeneous germline variants were removed and a copy number variation (CNV) analysis was performed to identify all aberrations and improve the filtering of germline variants. Genes were then compared to the AmygAc groups outcome to evaluate the influence with regard to the gene mutations. The group comparison was performed based on the level of genes and pathways. For the statistical results the coefficient of a binary logistic regression model with applied Wald Test was used. All results for each tested gene and pathway including the P-value (supplementary excel).

### Spatial Transcriptomics

For the RNA quality control of the 20 selected FFPE tissue blocks the commercially available Agilent’s Bioanalyzer assay was used. The RNA quality of the tissue was assessed by calculating DV200 of RNA extracted from freshly collected tissue sections. RNA was extracted using the RNeasy FFPE kit (Qiagen) following manufacturer’s instructions. Spatial transcriptomics (ST) was performed according to 10x Genomics protocols for FFPE tissue. For the analyses, a dedicated area of 6.5x 6.5 mm (capture area of the slides) containing tumor and non-tumor tissue was selected, half-cut and cut into 5 µm slides. The slides were transferred in a RNAse free water bath and placed on the capture area of the 10x Genomics slides, dried and post-processed by the Genomics Core Facility of Medical University of Vienna. First, a H&E staining was performed with subsequent imaging followed by hybridization, ligation and barcoding. Subsequently, a library was constructed and sequencing was performed. The capture of gene expression information for ST slides was performed by the Visium Spatial platform of 10x Genomics through the use of spatially barcoded mRNA-binding oligonucleotides in the default protocol. - Eighteen spatial transcriptomics slides were included in the study, comprising nine samples from the AmyAc-high group and nine samples from the AmygAc-low group. One sample (V11Y17-078_A1) was excluded further based on quality control assessment. This sample showed a markedly reduced library size and therefore a substantially lower contribution of tumor tissue compared to all other samples (see Supplementary Figures).

Raw gene count matrices in .H5AD format were obtained from SpaceRanger (version 4.0.1) aligned to GRCh38. Pathology assisted annotation of the spots in “tumor” and “healthy tissue” in each slide was performed using the Loupe Browser (version 9).

### Differential Gene Expression

Filtered feature matrices outputs of tumor tissue spots were aggregated per slide to generate pseudobulk expression profiles. Genes with zero counts across all samples were removed and lowly expressed genes were filtered by requiring a minimum count of 10 in at least 8 samples, corresponding to the smallest experimental group size. Library size normalization was performed using the DESeq2 [21] (version 1.46.0) *estimateSizeFactors* function with the *poscounts* option in R (version 4.4.3).

Differential gene expression between high-stress and low-stress conditions was assessed using DESeq2, fitting a negative binomial generalized linear model with stress status as the explanatory variable. Log2 fold changes were estimated and stabilized using *apeglm* shrinkage (version 1.28.0). Statistical significance was evaluated using Wald tests. P-values were adjusted for multiple testing using the Benjamini–Hochberg false discovery rate procedure.

### Gene Set Enrichment Analysis

Gene set enrichment analysis (GSEA) was performed using a preranked approach with gseapy (version1.1.11) in Python (version 3.9.23). All expressed genes derived from the differential expression analysis were ranked according to the DESeq2 Wald statistic. No additional fold-change or p-value cutoffs were applied prior to ranking, ensuring that the full transcriptome contributed to pathway-level inference.

Enrichment analysis was conducted using the Hallmark gene set collection from the Molecular Signatures Database (MSigDB)[21]. Gene sets containing fewer than 15 or more than 500 genes were excluded. Statistical significance was assessed using normalized enrichment scores and permutation-based p-values. Gene sets with a false discovery rate (FDR) < 0.25 were considered significantly enriched, in accordance with established GSEA recommendations[22].

### Cell Type Analysis

Cell-type enrichment analysis was performed in R (version 4.5.2) using the immunedeconv package[23](version 2.1.0) with the xCell method[24] (version 1.1.0). XCell is a single-sample GSEA-based method that estimates relative enrichment from bulk transcriptomic data. xCell enrichment scores were computed for all predefined cell-type signatures and interpreted as relative enrichment scores rather than absolute cell fractions. Differences in cell-type enrichment between high-stress and low-stress groups were assessed using Wilcoxon rank-sum tests on per-sample enrichment scores. P-values were adjusted for multiple testing using the Benjamini–Hochberg false discovery rate procedure.

### Multiplexing-IF

For multiplexing immunofluorescence (MP-IF), FFPE tissue blocks were cut in 3 µm slices and placed central on a coated microscope slide. Tissue sections of the same 20 patients as for ST were stained. For MP staining, six different antibodies were used (PD-L1 Optibody, CD8, CD3, PD-1, CD45RO, CK-pan-Cytokeratin and 4′,6-diamidino-2-phenylindole (DAPI)) in an autostainer system Bond RX (Leica Biosystems Inc., Vienna Austria). Assessment was performed with multispectral imaging using the Vectra (Akoya Biosystems, Marlborough, US, software version 3.0) separately for each color channel/antibody, and for post processing for multispectral images HALO Image Analysis Platform (Version 3.4. Albuquerque, NM: Indica Labs, Inc.; 2021) was used. Cell-based features were extracted from whole slide images (WSI) cut into tiles of 1,700µm, tiles with more than 100 cells were kept with 35 to 220 tiles/WSI. Results from all tiles were aggregated to represent WSI correctly. The resulting fifteen phenotypes (supp table 2) were analyzed with regard to total tissue, tumor area, TME, and stress groups using multispectral imaging post-processed in HALO and quantified as number of cells expressed in % of total cells. Additionally, cells within a radius (15, 30, 50 µm) representing the number of cells from one phenotype within a radius of another were explored.

Phenotype features from the tissue samples were analyzed using in a first step MANOVA to assess multiple dependent variables, and to test for influence of independent variables as the histology. For the significant results, a one-way ANOVA was calculated to dissect the MANOVA results. To exclude a bias for features which are not normally distributed and to test for any differences in the groups, the non-parametric Kruskal-Wallis test was used.

Three key immune markers were investigated: the overall proportion of cytotoxic T lymphocytes (CTL %), and the number of CTLs in proximity to PD-L1-expressing cells as well as to PD-1-expressing cells. Both parameters had strongly non-normal and right-skewed distributions and unequal variances, as usually observed in spatial immune cell data and small sample sizes. To meet the assumptions of parametric testing and to ensure robustness against outliers, values are reported as medians and compared using two-sided Mann–Whitney U tests with Benjamini–Hochberg correction across the predefined marker family.

To assess the impact of higher AmygAc on the immune landscape of the TIME, multiplexing immunofluorescence (MP-IF) was performed, and among all six antibodies and resulting 15 phenotypes (see supplemental Table 1), we focused on a predefined cytotoxic immune-checkpoint marker family comprising global cytotoxic T-cell infiltration (CTL %) and spatial CTL proximity to PD-L1/CD274- and PD-1-expressing cells (#CTL around PD-L1, #CTL around PD-1), as these markers jointly capture cytotoxic immune abundance and spatial engagement of CTLs with the PD-1/PD-L1 inhibitory checkpoint axis.

For PD-L1 scores for the immunotherapy cohort and skewed distributions, markers are reported as medians and compared using two-sided Mann–Whitney U tests with Benjamini– Hochberg correction. Global CTL infiltration was numerically lower in AmygAc-low patients but did not reach significance according to the Mann-Whitney U test (with Benjamini-Hochberg correction P = 0.09).

## Results

The final study cohort (Figure 1) consisted of 256 treatment-naïve lung cancer patients from four different centers (median age 65 years, 50% male). Most patients presented with stage III–IV disease (193 cases) and adenocarcinoma being the predominant histological subtype.

### Inflammatory markers fail to predict outcome

From the [^18^F]FDG PET images, beside the metabolic activities of the amygdala (AmygAc) also bone marrow as marker of inflammatory activation was extracted, see Figure 2 for representative images. Baseline inflammatory markers, including CRP and leukocyte count, were markedly elevated indicating an active smoking habit in the vast majority of patients. However, they didn’t not differ between patients of the survival groups (4.6 vs. 3.8 mg/dl (total cohort 4.4 mg/dl) (Table 1 and Supplement for more details).

Total lesion glycolysis and amygdala metabolism are associated with survival While PET and blood-based inflammatory parameters did not show significant predictive potential, lower AmygAc and total lesion glycolysis (TLG) as well as surgical treatment were positively associated with patients’ one-year survival (all P < 0.001), The mean follow-up time for the entire cohort was 27 months (range 0.3-84). Among survivors (censored patients), the mean follow-up time from baseline to last contact was 40 months (range 13-84), whereas patients who died were followed for a mean of 16 months until death.

Patients who did not receive neo-adjuvant ICI therapy had either radio- or chemo-therapy or surgery or, in case of 27 patients, a combination of more than one of those.

Metabolic amygdala activation predicts resistance to immune checkpoint inhibitor therapy AmygAc was considered an objective central and host-derived biomarker retrospectively assessed from 79 therapy naïve diagnostic scans of lung cancer patients who subsequently received neoadjuvant ICI therapy. Response to therapy was assessed based to the RECIST criteria from the tumor volume change determined morphologically from CT performed on average (146 ± 197) days after the baseline scan. This cohort was dichotomized into 34 responders (i.e. having a tumor volume reduction of at least 30 %) with a mean survival of 26 months and 45 non-responders with a mean survival of 14 months. Amongst responders, the AmygAc was significantly lower compared to the sub-group of non-responders (0.76 versus 0.81, P= 0.003 (student’s t-test), see also Figure 3a). This was still highly significant in a logistic regression model after multi-variate adjusting for survival (P = 0.004), leading to a sensitivity of 0.59 and a specificity of 0.84 (Figure 3b). Thus, assessing AmygAc at baseline provides a robust prognostic marker for ICI-resistance.

**Figure 3:**
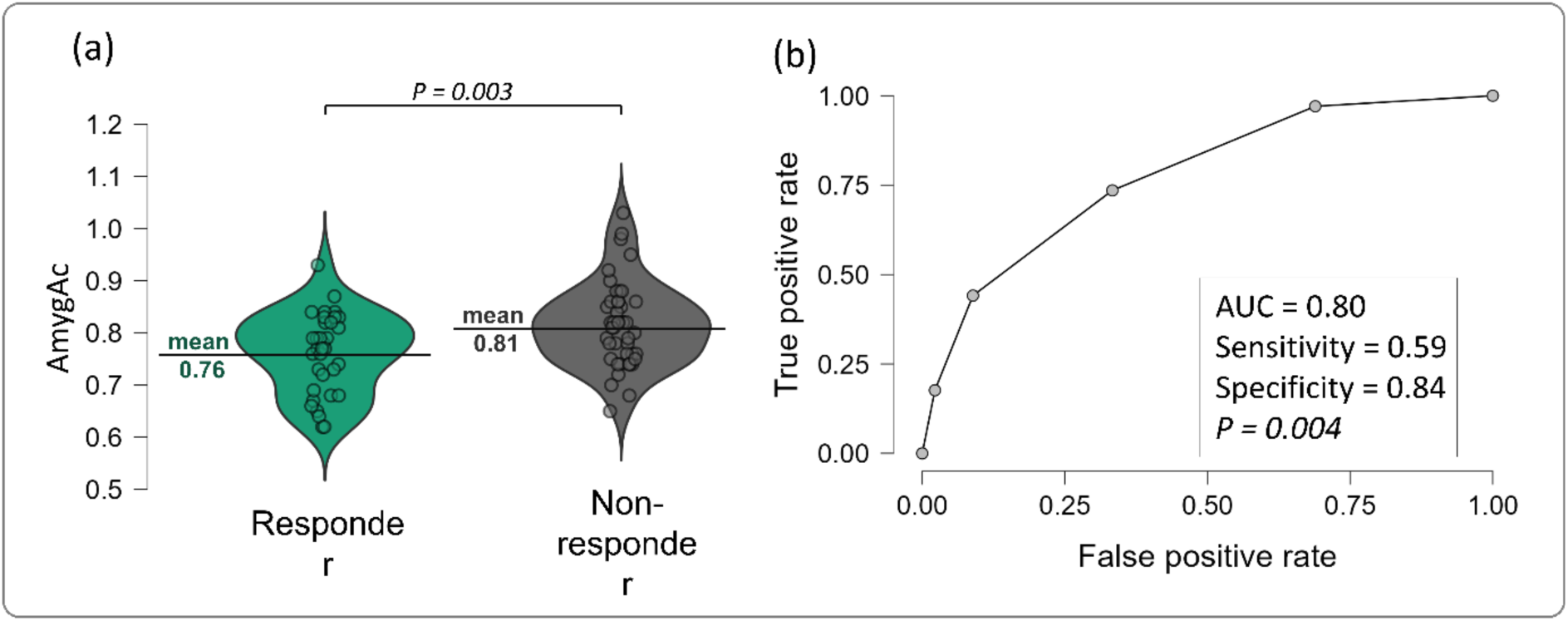
(a) Violin plot of baseline amygdala activity from 79 patients (45 non-responder and 34 responder) receiving neoadjuvant ICI therapy; (b) Receiver Operating Characteristic (ROC) curve showing the discriminative performance of baseline amygdala activity (AmygAc)in distinguishing responder from non-responder, with an area under the curve (AUC) of 0.8.

Amygdala activation affects outcome directly and via immunomodulatory processes After proving the association of PET AmygAc with ICI therapy response, we evaluated baseline parameters for outcome prediction and used surgery specimens from therapy naïve tumor material for an in-depth molecular analysis to relate outcome parameters with molecular profiles and response assessment. First, a Cox-regression analysis revealed that tumor activity (TLG) and tumor stages had a significant impact on the one-year survival (both P-values < 0.001, Figure 4a), as well as AmygAc (P = 0.001), while other clinical parameters (C-reactive protein, leucocytes, age, sex, body mass index BMI, bone marrow activity) were not significant. AmygAc remained a robust predictor for one-year survival after multivariate adjustments (see Figure 4a). To investigate overall survival, AmygAc was dichotomized using a survival-ROC analysis into low (signal-to-noise ratio lower than 0.86, for details see supplement) and high AmygAc, associated with low and high chronic stress exposure, respectively. A Kaplan-Meier analysis revealed a significantly (log-rank test P = 0.003) longer survival in patients with low AmygAc exposure, identifying AmygAc as an imaging biomarker for poor outcome of similar predictive power as TLG (see Figure 4c), which was so far the only available image-derived biomarker for survival.

**Figure 4:**
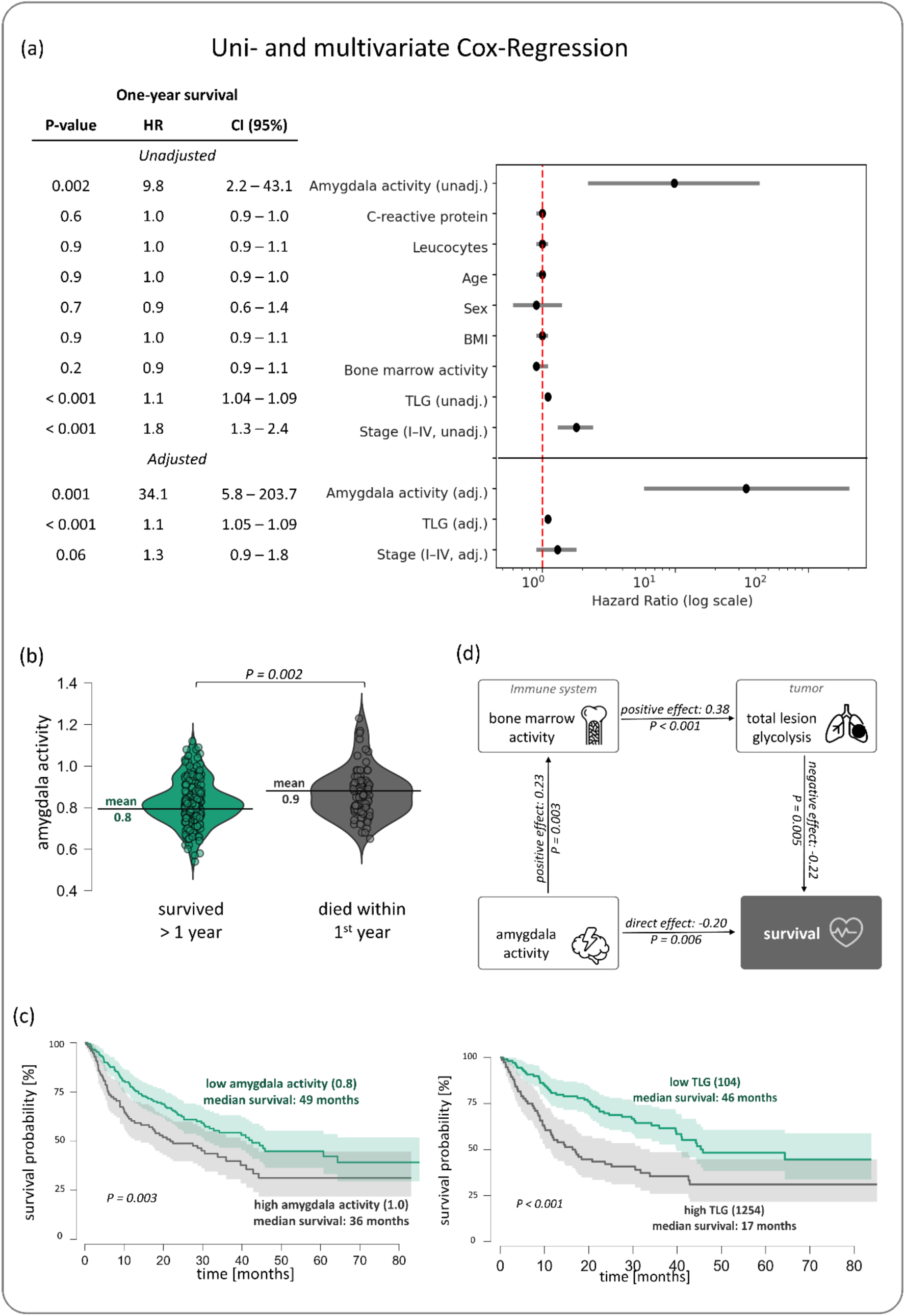
(a) Cox-regression analysis of the impact of all available parameters on the one-year survival including a forest plot of the hazard ratio (HR) showing the 95 % confidence interval (CI). (b) Violin plot of the left amygdala activity in the group which survived the first year after baseline scan (green) and which died within the first year (gray), with the P-value from the student’s t-test. (c) Kaplan-Meier analyses of the overall survival of the group with low amygdala activity associated with low chronic stress exposure at baseline (green) and with high chronic stress (gray) with the P-value from the long rank test, and in comparison, to the groups with low and high tumor marker (TLG: total lesion glycolysis). (d) Mediation analysis showing that there is significant direct effect of the left amygdala activity on the one-year-survival, but also an indirect effect mediated by the marker for the inflammatory activation (bone marrow activity), influencing the tumor marker (total lesion glycolysis).

These results robustly show that AmygAc not only impairs ICI susceptibility, but leads to lower survival after lung cancer diagnosis, independently of inflammatory state, tumor burden or treatment modality. We performed a mediation analysis (Figure 4d, for details see also supplement), allowing to test whether the effect of high AmygaAc on the one-year-survival was mediated through other acquired parameters and thus to assess causality. We detect a robust direct effect of AmygAc and Survival (−0.2, P = 0.006) and a significant overall indirect effect (0.5, P < 0.05, not shown), showing that the relation was serially mediated via the bone marrow activity and the tumor activity TLG.

Multiplexing and transcriptomics confirm AmygAc-dependent alterations of TIME To identify the amygdala mediated signals affecting patient outcome, we aimed to molecularly characterize the therapy naïve TIME of patients with high and low AmygAc. We acquired tumor tissue samples from patients which underwent surgery before any other therapeutic intervention. The cohort consists of 10 patients with high and 10 with low AmygAc at baseline. Patient selection was based on the availability of diagnostic pathology material and balanced representation of gender and histology (AC vs SCC). In the high AmygAc group overall survival was 31 ± 17 months, compared to 54 ± 21 months (P = 0.01) in patients with low AmygAc. To exclude any inadvertent genetic confounders driving the differences, we analyzed oncogenic drivers. We especially focused on the most common lung cancer mutations (P53, EGFR, HER2, KRAS, BRAF, ROS1, ALK, PI3KCA, MET, FGFR1, DDR2, RET, NTRK, MEK1) and performed a comparative group analysis, which revealed no statistically significant differences in the mutational profile of the tumors or in pathway analyses (for details, see supplement excel tables 1 and 2).

We confirmed the histologically diagnosis and the tumor and TIME was delineated by pathologists before multiplexing and spatial transcriptomics analysis (see Figure 5b). Quality control of the RNA was sufficient with median coverage of ∼ 200x using a kit, however one sample was excluded from further downstream analysis after showing decreased library size (see also supplement). Hence, integrated transcriptome and pathway analysis between the high- and low-AmygAc group reveals decreased interferon-γ (INF-γ) signaling, with a negative enrichment score (NES) of −1.65 with a nominal p-value < 0.001 and a false discovery rate (FDR) of 0.015. The negative NES indicates that genes transcriptionally activated by INF-γ signaling are preferentially enriched among genes upregulated in the lowAmygAc group, suggesting attenuated INF-γ pathway activity under neuronal activity in the amygdala region (Figure 5e). Top five-ranked up- and downregulated pathways (supplemental Figure 4) are associated with increased expression of proliferation pathways and decreased expression of inflammatory and metabolic markers suggesting potential lowered immune response, muscle wasting and altered metabolism in the high AmygAc group.

**Figure 5:**
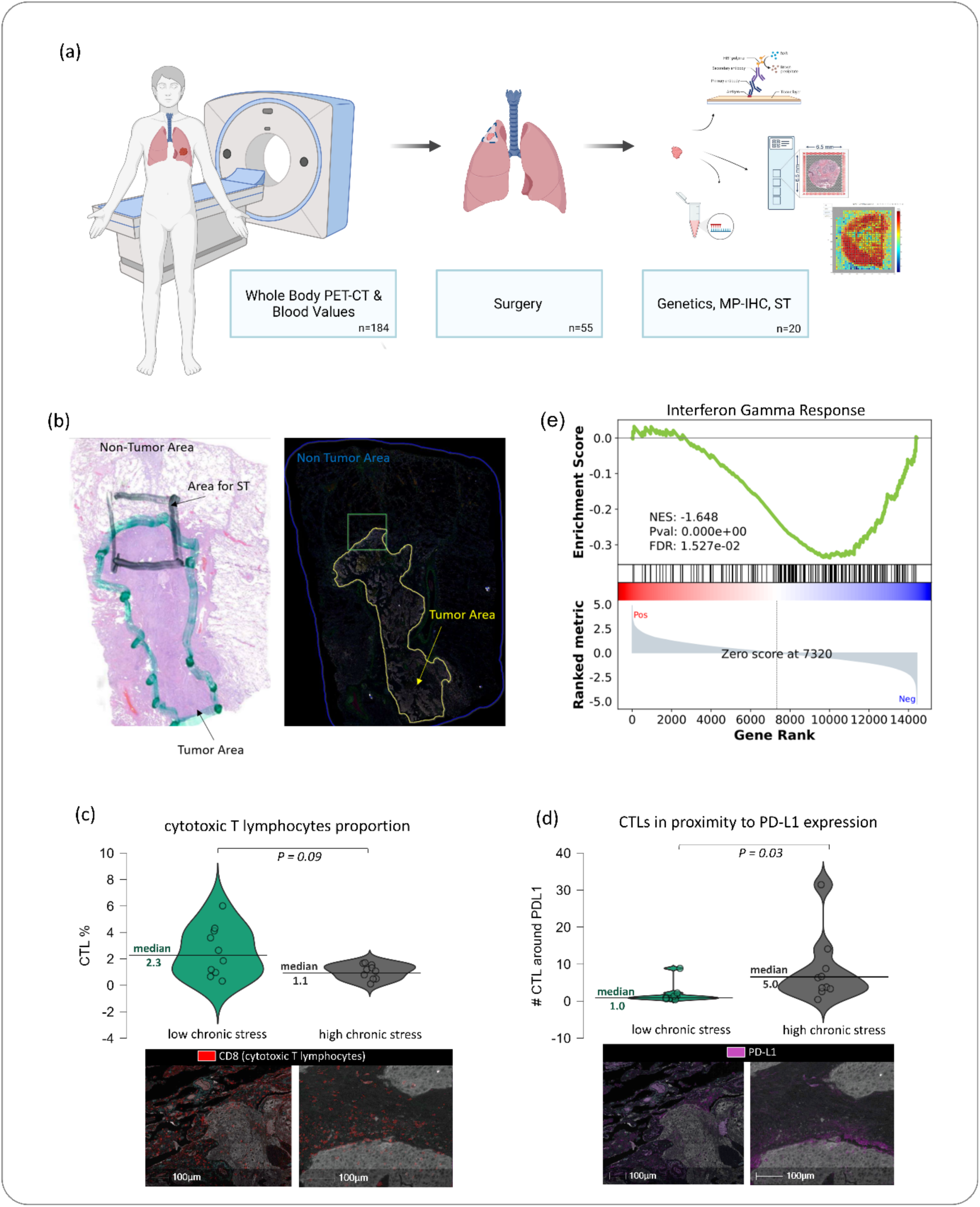
(a) Illustration of General Hospital cohort workflow including the biological samples. N=55 patients underwent surgery of which 20 tumor blocks (10 of each group) were taken for OMICS analyses. (b) Representative H&E (left) and multiplexing images (right) with annotations: Square area indicates the area taken for Visium 10x Genomics (spatial transcriptomics); green area (H&E, segmentation by a pathologist) and yellow area (multiplexing immune fluorescence, MP-IF) represents the tumor area (additionally indicated with an arrow). Surrounding tissue is indicated as non-tumor area with an arrow. (c) Violin plots of the relative expression of cytotoxic T lymphocytes (CD8+/CD3+) on total tumor block expressed in percent of patients with low and high left amygdala activity at baseline (representative staining below). (d) Violin plots of cytotoxic T lymphocytes (CD8+/CD3+) in proximity to Programmed Death-Ligand 1 (PD-L1) (radius analysis) between patients with low and high left amygdala activity at baseline (representative staining below). P-values from two-sided Mann–Whitney U tests are Bonferroni corrected for multiple testing. (e) Gene set enrichment analysis (GSEA) of the Hallmark Interferon gamma response pathway. The pathway is significantly negatively enriched in high-stress samples (NES = −1.65, FDR q = 0.015), indicating suppression of interferon-γ–related immune signaling relative to low-stress samples. Abbreviation: CD8+/CD3+ cytotoxic T lymphocytes (CTL); Programmed Death-Ligand 1 (PD-L1)

Additionally, xCell analysis showed tendencies of differences in the enrichment of multiple immune cell signatures (supplemental Figure 4 and 5).

To assess the impact of high AmygAc on the immune landscape of the TIME, multiplexing immunofluorescence (MP-IF) was performed, and among all six antibodies and resulting 15 phenotypes (see supplemental Table 1), we focused on a predefined cytotoxic immune-checkpoint marker family comprising global cytotoxic T-cell infiltration (CTL %) and spatial CTL proximity to PD-L1/CD274- and PD-1-expressing cells (#CTL around PD-L1, #CTL around PD-1), as these markers jointly capture cytotoxic immune abundance and spatial engagement of CTLs with the PD-1/PD-L1 inhibitory checkpoint axis. Given small group sizes and skewed distributions, markers are reported as medians and compared using two-sided Mann–Whitney U tests with Benjamini–Hochberg correction. Global CTL infiltration was numerically lower in patients with higher AmygAc but did not reach significance according to the Mann-Whitney U test (with Benjamini-Hochberg correction P = 0.09). In contrast, CTL proximity to PD-L1/CD274-expressing cells was significantly increased in those patients (#CTL around PD-L1: median 5.04 vs. 1.02; P 0.03), whereas CTL proximity to PD-1-expressing cells showed a non-significant trend in the same direction (#CTL around PD-1: median 19.76 vs. 6.06; P = 0.09). This suggests a coordinated shift toward reduced PD-L1-associated CTL clustering under low AmygAc.

We further evaluated if the engagement between PD-L1 expressing cells and CTL is driven by higher PD-1 or PD-L1 expression and showed the independency as neither PD-1 nor PD-L1 expression differ between the groups (see supplement figure 6).

## Discussion

Our findings provides additional support to the fact that routine whole-body [18F]FDG PET simultaneously captures tumor- and host-derived imaging biomarkers, which can provide clinically relevant information beyond lesion-centered metabolic assessment. Amygdala metabolic activity (AmygAc) independently predicted survival and response to immune checkpoint inhibition and demonstrated predictive performance comparable to total lesion glycolysis (TLG), an established marker of metabolically active tumor burden widely used in clinical management [25]. Current biomarkers of immunotherapy response primarily rely on tumor-intrinsic characteristics such as PD-L1 expression, tumor mutational burden and IFN-γ-related gene signatures, all of which are influenced by spatial heterogeneity and methodological variability [26,27]. Our findings suggest that host-derived imaging biomarkers obtained from routine whole-body [^18^F]FDG PET provide complementary biological information that is not captured by established tumor-based biomarkers. Although elevated AmygAc has previously been associated with chronic psychological stress, depression, anxiety and adverse clinical outcomes, the biological mechanisms underlying its prognostic significance for diseases of the periphery like cancer have remained largely undefined. Chronic psychological stress is a complex and multifactorial process, and unhealthy coping behaviors may also contribute indirectly to adverse clinical outcomes [28,29]. Rather than directly measuring psychological stress, our study identifies AmygAc as an objective host-derived imaging biomarker associated with distinct alterations of the tumor immune microenvironment and reduced susceptibility to immune checkpoint inhibition. Unlike questionnaire-based assessments, AmygAc can be quantified retrospectively from routinely acquired [^18^F]FDG PET examinations without additional imaging or patient burden [9,11,30–32].

These findings further extend the role of [18F]FDG PET beyond assessment of tumor metabolism alone and suggest that simultaneous characterization of host and tumor metabolism may improve prognostic assessment in NSCLC. Systemic inflammatory and immune biomarkers have repeatedly been associated with poor ICI outcomes [33]. Current biomarker strategies have primarily focused on systemic inflammation and T-cell dysfunction [34–36], but their variability and lack of specificity limit their clinical predictive value. It is therefore important to note that patients with NSCLC generally display strongly elevated inflammatory markers including CRP and LEUCOCYTES due to smoking-related comorbidities, including chronic obstructive pulmonary disease, and therefore incompletely capture the host biology relevant for treatment susceptibility [37,38]. NSCLC therefore provides an interesting paradigm where the influence of the inflammatory burden may not confound the analysis of a stress-related influence on the TIME. Our data suggest that AmygAc correlates with stress-related biological processes that are distinct from systemic inflammatory pathways and provide additional prognostic information beyond tumor metabolic burden and established inflammatory biomarkers. The host-derived neural imaging biomarker AmygAc provides a robust measure capturing downstream biological processes contributing to immunesuppression in the TIME independent of systemic inflammatory burden.

These AmyAc-dependent immunesuppression in TIME was associated with reduced IFN-γ release and lower cytotoxic T-cell infiltration based on our analysis of 20 treatment-naïve tumors. Transcriptomic pathway analysis identified IFN-γ target programs as consistently downregulated in patients with elevated AmygAc; additional inflammatory pathways (IFN-α and TNF-α via NF-κB) [39,40] were likewise attenuated, whereas proliferation-related programs were enriched. Multiplex immunohistochemistry further demonstrated increased spatial engagement between CD8+ T cells and PD-L1-expressing cells in tumors from patients with elevated AmygAc, but no differences in PD-L1 expression in either the treatment-naïve or immunotherapy cohorts were observed. It is important to note that TLG did not differ between the high and AmygAc patients. Collectively, these findings indicate that elevated AmygAc is associated with an immunologically suppressed TIME characterized by impaired IFN-γ signaling and reduced cytotoxic immune activity, which cannot be explained by TLG. Although IFN-γ is a central regulator of antitumor immunity in lung cancer through activation of effector lymphocytes, previous reports have differed regarding the relationship between IFN-γ signaling, PD-L1 expression and CD8+ T-cell infiltration in resistant disease[41]. While we observed IFN-γ expression was consistently reduced in patients with elevated AmygAc, so far there are only isolated studies linking chronic psychological stress with reduced IFN-γ signaling [42,43].

Importantly, AmygAc is derived from the brain component of routinely acquired whole-body [18F]FDG PET examinations and therefore requires neither additional tracer administration nor dedicated neuroimaging protocols. This demonstrates how a single imaging examination can simultaneously characterize tumor metabolism and host biology, supporting a systems-level approach to [18F]FDG PET that extends beyond conventional lesion-based assessment. Therefore, AmygAc constitutes a strong candidate for the first biomarker reflecting the influence of the host status on therapy response and might overcome current limitations in clinical stratification pipelines, which focus on tumour specific parameters but are plagued by significant heterogeneity in therapy response.

Several limitations should be acknowledged. The retrospective design precludes causal inference, and AmygAc should therefore be interpreted as an imaging biomarker associated with, rather than a direct measure of, chronic psychological stress. Furthermore, objective measures of stress, including psychometric assessments or circulating biomarkers such as cortisol, were not available. Prospective studies integrating molecular profiling with validated physiological and psychological measures will be important to further define the biological basis of elevated AmygAc and its relationship to antitumor immunity [10,42,44–46].

In conclusion, our findings identify AmygAc as a readily accessible host-derived imaging biomarker, which are currently lacking from stratification approaches. AmygAc may thus complement established tumor PET biomarkers for prognostic assessment and prediction of immunotherapy response. Prospective validation is warranted to determine its clinical utility for patient stratification and to evaluate whether interventions targeting the biological pathways associated with elevated AmygAc can improve immunotherapy efficacy. More broadly, our results support the concept that whole-body [18F]FDG PET can simultaneously characterize host and tumor biology, providing mechanistic insights into immune regulation that extend beyond conventional assessment of tumor metabolism alone.

## Supporting information

Supplementary Metadata and Omics Data

## Abbreviations

AC: adenocarcinoma
ADSQ: adenosquamous carcinoma
AKH: Vienna General Hospital (Allgemeines Krankenhaus Wien)
AmygAc: amygdala metabolic activity
ANOVA: analysis of variance
AUC: area under the receiver operating characteristic curve
BMI: body mass index
CI: confidence interval
CNV: copy number variation
CRP: C-reactive protein
CT: computed tomography
CTL: cytotoxic T lymphocyte
DAPI: 4′,6-diamidino-2-phenylindole
DV200: percentage of RNA fragments longer than 200 nucleotides
FDG: fluorodeoxyglucose
FDR: false discovery rate
FFPE: formalin-fixed paraffin-embedded
fMRI: functional magnetic resonance imaging
GSEA: gene set enrichment analysis
H&E: hematoxylin and eosin
HR: hazard ratio
ICI: immune checkpoint inhibitor
IF: immunofluorescence
IFN-α: interferon alpha
IFN-γ: interferon gamma
LAFOV: long axial field of view
LCC: large cell carcinoma
MANOVA: multivariate analysis of variance
MP-IF: multiplex immunofluorescence
MSigDB: Molecular Signatures Database
MTV: metabolic tumor volume
NES: normalized enrichment score
NSCLC: non-small cell lung cancer
OSEM: ordered-subset expectation maximization
PD-1: programmed cell death protein 1
PD-L1: programmed death-ligand 1
PET: positron emission tomography
PSF: point spread function
RECIST: Response Evaluation Criteria in Solid Tumors
ROC: receiver operating characteristic
SCC: squamous cell carcinoma
SNR: signal-to-noise ratio
SNRmax: maximum signal-to-noise ratio
SNRmean: mean signal-to-noise ratio
ST: spatial transcriptomics
SUV: standardized uptake value
SUVmax: maximum standardized uptake value
SUVmean: mean standardized uptake value
TBR: target-to-background ratio
TIME: tumor immune microenvironment
TLG: total lesion glycolysis
TOF: time of flight
VOI: volume of interest
WBC: white blood cell count
WSI: whole-slide image

## Acknowledgements

We kindly acknowledge our colleagues Elisabeth Gurndorfer (Pathology) and Sophie Derdak, Christoph Friedl and Martin Bilban at the Genomics Core Facility of Medical University in Vienna for their valuable contributions to the spatial transcriptomics. We acknowledge CF Prot of CIISB, Instruct-CZ Centre, supported by MEYSCR (LM2023042)) and European Regional Development Fund-Project „UP CIISB“ (No. CZ.02.1.01/0.0/0.0/18_046/0015974) and, specifically Vojtech Bystry and Boris Tichy for sequencing and analysis of the tumor markers. Additionally, we thank Lucian Beer and Daria Kifjak from Klinik Floridsdorf for their support and collaboration. CV acknowledges financial support for the salary of CTSM by the Austrian Science Fund through FW771G0201 “EPILUCAFS” Grant-DOI 10.55776/PIN5320023. We further like to cite and acknowledge BioRender.com as we used the tool for creating figure 5a.

## Authors Contributions

B.K.G. and C.V. contributed equally to this work. B.K.G. and C.V. performed the formal analyses, developed the methodology, generated the visualizations and wrote the original draft of the manuscript. C.T.S.M., L.B., A.F., L.H., N.H., S.H., L.K., D.K., K.K., O.C.K., C.L., A.L., T.S.N., F.O., R.K., O.S., B.S., C.A., K.T., M.T., H.W. and J.Y. contributed to patient recruitment, data acquisition and investigation. O.C.K., H.P., D.T. and J.Y. curated the datasets. C.S. performed software development, validation and contributed to formal analysis. T.S. contributed to methodological development. S.G. and M.H. conceived and supervised the study. S.G. contributed to conceptualization, drafting of the manuscript and manuscript revision. M.H. contributed to conceptualization, methodological design and manuscript revision.

## Data availability

The imaging and clinical datasets generated and analyzed during the current study are not publicly available due to their large size and patient privacy considerations but are available from the corresponding author upon reasonable request.

