## Supplementary Information for "Amygdala metabolic activity on [^18^F]FDG PET is associated with survival, response to immune checkpoint inhibition, and tumor immune signaling in non-small cell lung cancer"

**Supplement**

**Imaging data**

The retrospective cohort of therapy-naïve lung cancer patients consisted of contributions of four centers: 174 subjects from the General Hospital in Vienna, Austria (AKH), 19 subjects from Klinik Floridsdorf in Vienna, Austria, 5 from Universitätsklinikum Augsburg, Germany, and 58 from University of Leipzig, Germany. From the subjects from AKH, 20 tissue samples were chosen for further analysis, and 38 follow-up CT scans were available after therapy allowing to evaluate response from morphological changes. From the other three centers (Leipzig, Augsburg and Floridsdorf), altogether 79 follow-up CT scans after ICI therapy were available.

In the images, for the amygdala activity, a cubic volume of interest (VOI) was placed in the left amygdala, mean and maximum tracer concentrations were measured; mean brain background activity was determined from the temporal lobe. The left amygdala activities were normalized to the temporal lobe background, thus the signal-to-noise-ratios SNR_mean_ and SNR_max_ served as primary and secondary markers for chronic stress, whereas the left amygdala SNR_mean_ was denominated as AmygAc. Tumor volumes were delineated semi-automatically using liver segment VIII as reference region, mean tumor volume (MTV) and tumor mean standardized uptake value (SUV_mean_) was measured and the total lesion glycolysis (TLG) was further used as a marker of tumor metabolic activity. [^18^F]FDG maximum uptakes in the bone marrow and the adrenal glands were normalized to blood pool activity. Blood pool activity was obtained through a representative intravascular VOI in the ascending aorta. The uptake of the adrenal glands and of the brain background were used to evaluate imaging differences between the participating centers.

**Mediation analysis**

Mediation analysis was performed to examine whether the association between AmygAc and survival was transmitted through downstream biological processes. Mediation analysis decomposes the total effect of an exposure (AmygAc) on an outcome (survival) into a direct effect and indirect effects operating through intermediate variables (mediators). Using linear regression with bootstrapping, AmygAc was modeled as the independent variable, survival as the dependent variable, and bone marrow metabolic activity and TLG as serial mediators. This approach tests whether chronic stress (AmygAc) influences survival partially by modulating systemic immune activation (bone marrow activity), which in turn affects tumor metabolic burden (TLG).

**Results**

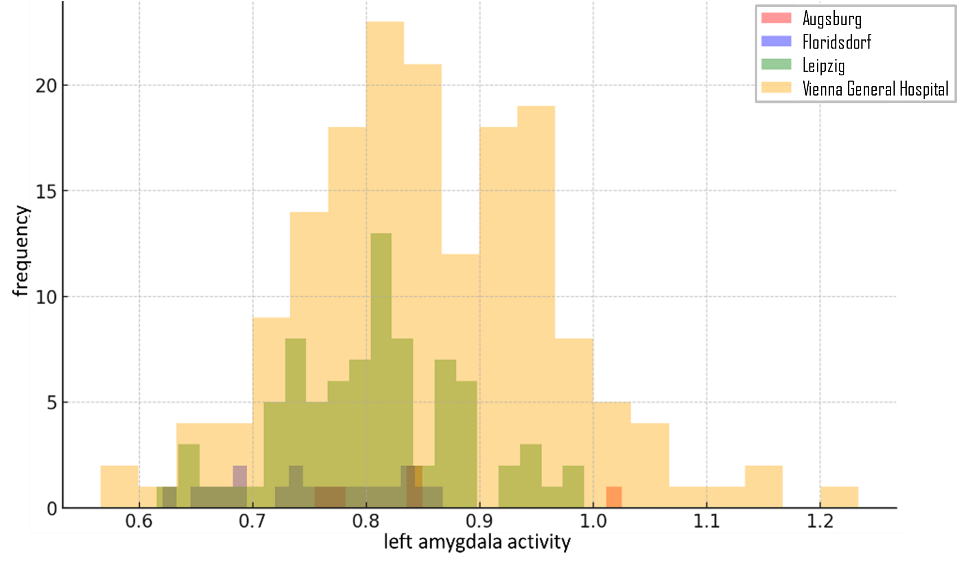
To explore systemic differences between the participating centers, regarding imaging parameters adrenal glands and brain background were analyzed with student’s t-tests, i.e. areas were investigated which might be less affected by tumor, inflammation or health status. No significant difference was found between all centers neither in brain background (all P-values > 0.2) nor in adrenal glands (P-values > 0.8). However, in particular between the both major cohorts, Vienna General Hospital and Leipzig, major differences were found in clinical parameters, with the Vienna General Hospital cohort being significantly older (66 versus 60 years, P < 0.001), having a longer survival (31 versus 16 months, P < 0.001), a lower tumor stage (median III versus median IV) and less non-responder (20 versus 44, P = 0.008). The image-derived markers for chronic stress and immune response, left amygdala and bone marrow activity, respectively, were significantly higher (0.86 versus 0.81, P < 0.001, and 1.36 versus 1.15, P = 0.02), see also Figure 1.

Supplementary Figure 1: overlaying histograms of the primary chronic stress marker according to the different participating centers.

A survival-ROC analysis was performed to estimate the optimum amygdala activity threshold for the one-year survival, leading to a threshold 0.86, i.e. stratifying patients with higher values into those with exposure to chronic stress. This analysis was also performed with the secondary stress marker, the maximum amygdala activity, resulting in a threshold of 1.08, and with the TLG, resulting in a threshold of 320.

Regarding one-year survival, both markers for chronic stress, mean and maximum left amygdala activity, were significantly higher in those who died within the first year: 0.87 versus 0.82 (P = 0.002) in case of mean and 1.14 versus 0.73 (P = 0.006) in case of maximum amygdala activity.

Note that we also investigated right amygdala activity, however, results regarding survival were comparable, but less or not significant (P values around 0.08), thus the right amygdala was excluded from detailed and further analysis. The Cox regression for the maximum amygdala was also significant with a hazard ratio of 3.6 (95% CI: 1.13 – 11.4, P = 0.03) and also the overall survival in a Kaplan-Meier analysis and a log-rank test with 51 months survival in those with low and 39 months with high maximum amygdala activity (P = 0.01). A Kaplan-Meier analysis of the TLG resulted in a significantly longer survival of patients with low TLG (< 320) of 52 months (versus 35 months, P < 0.001).

A weak but significant positive correlation of R = 0.2 was found between amygdala and bone marrow activity (P = 0.001 for mean and P = 0.01 for maximum amygdala activity), and between bone marrow activity and CRP (R = 0.3, P < 0.001), suggesting a relation between inflammatory reaction and chronic stress. Negative correlations with survival were found in case of amygdala activity (R = -0.2, P = 0.001 and P = 0.003 with mean and max, respectively) and in case of bone marrow activity (R = -0.2, P = 0.009), showing that higher chronic stress and higher inflammatory activity are related to a shorter survival. Survival was also negatively correlated with TLG: R = -0.25, P < 0.001, and positively with bone marrow activity: R = 0.3, P < 0.001. The relationship between these markers were in detail analyzed with a mediation analysis, allowing to assess causal connections between correlating variables. A triangular relationship was not significant from amygdala as predictor and survival as outcome, neither with TLG as mediator (indirect effect P = 0.3), nor with bone marrow activity (indirect effect P = 0.1), nor with CRP (indirect effect P = 0.6), the multiple model with two mediators involved was highly significant, with bone marrow as first and TLG as second mediator (indirect effect P < 0.05). This suggests a serial impact from chronic stress on inflammatory processes, which in turn than affect the aggressivity of the tumor.

**Missing values**

Age, BMI and CRP were missing from 3 %, 31 % and 25 % of all subjects, leucocytes were only available from the AKH cohort with 3 % missing values. Tumor-related histology findings were missing from 18 %, survival status from 2 %, TLG from 31 %. While all amygdala and brain background SUVs were available, 14 % of the adrenal gland and 3 % of the bone marrow SUVs were missing.

**Biological Data**

**Cell Type Analysis**

Cell-type enrichment analysis was performed in R (version 4.5.2) using the immunedeconv package(*5*) (version 2.1.0) with the xCell method (*6*)(version 1.1.0). XCell is a single-sample GSEA-based method that estimates relative enrichment from bulk transcriptomic data. xCell enrichment scores were computed for all predefined cell-type signatures and interpreted as relative enrichment scores rather than absolute cell fractions. Differences in cell-type enrichment between high-stress and low-stress groups were assessed using Wilcoxon rank-sum tests on per-sample enrichment scores. P-values were adjusted for multiple testing using the Benjamini–Hochberg false discovery rate procedure.

**MP-IF**

*Cell Markers used for multiplexing immunofluorescence staining*

| **Cell Type/**  **Phenotype** | **Cell Markers** | | | | | |
| --- | --- | --- | --- | --- | --- | --- |
|  | **CD3** | **CD8** | **CD45Ro** | **PD-1** | **PD-L1** | **CK** |
| T cell general | x |  |  |  |  |  |
| Cytotoxic T cells (CTL) | x | x |  |  |  |  |
| CTL memory | x | x | x |  |  |  |
| PD-1 Expression |  |  |  | x |  |  |
| PD-1 Expression on T cells | x |  |  | x |  |  |
| PD-1 Expression on CTL | x | x |  | x |  |  |
| PD-1 Expression on tumor cells (CK+) |  |  |  | x |  | x |
| PD-L1 Expression |  |  |  |  | x |  |
| PD-L1 Expression on T cells | x |  |  |  | x |  |
| PD-L1 Expression on CTL | x | x |  |  | x |  |
| PD-L1 Expression on tumor cells (CK+) |  |  |  |  | x | x |
| PD-1 + PD-L1 Expression |  |  |  | x | x |  |
| PD-1 + PD-L1 Expression on T cells | x |  |  | x | x |  |
| PD-1 + PD-L1 Expression on CTL | x | x |  | x | x |  |
| PD-1 + PD-L1 Expression on tumor cells (CK+) |  |  |  | x | x | x |

**Supplement table 1:** List of analyzed phenotypes according to the six stained immune markers for *multiplexing immunofluorescence staining* (MP-IF). Used abbreviations are CK, CK-pan-Cytokeratin; PD-1, programmed cell death protein 1 and PD-L1, programmed death-ligand 1.

For pathologist’s PD-L1 score, the staining platform Ventana Benchmark Ultra was used with either PD-L1-Klon 22C3 (Dako) or PD-L1 antibody 28-8 (abcam) (*7*). Combined score (CPS) was calculated using the number of PD-L1 stained cells (tumor cells, lymphocytes and macrophages) divided by the total number of viable tumor cells and multiplied with 100%. To meet the assumptions of parametric testing and to ensure robustness against outliers, values are reported as medians and compared using two-sided Mann–Whitney U tests with Benjamini–Hochberg correction across the predefined marker family.

**Results**

**Transcriptional analysis**

RNA quality control showed nucleic acid concentrations ranging between 160and 1350 ng/µL using Agilent’s Bioanalyzer assay.

Library size per sample, measured as the total number of RNA-seq counts derived from tumor tissue, shown on a log₁₀ scale. Sample V11Y17-078_A1 exhibited a markedly reduced library size (199,561 total counts) compared with all other samples and was therefore excluded from downstream analyses.

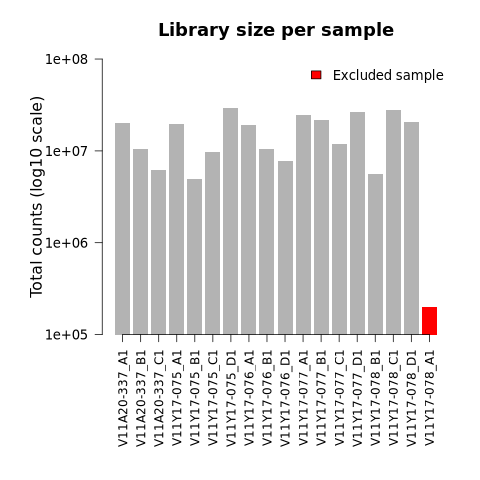

Supplementary Figure 2: Library size per sample, measured as the total number of RNA-seq counts derived from tumor tissue, shown on a log₁₀ scale.

**Pathway Enrichment Analysis**

The Pathway Enrichment Analysis between the high-stress and low-stress group reveals a negative enrichment score (NES) for IFN-γ and IFN-α response. Additionally, hallmarks of myogenesis, genes regulated by Nuclear Factor kappa-light-chain-enhancer of activated B-cells (NF-κB) in response to tumor necrose factor alpha (TNF-α) as well as early estrogen response are downregulated. Indicating potential, muscle wasting in stressed-patients, inflammatory and metabolic changes and altered stress response, as well as reduced ER-mediated transcriptional activity. While the upregulated top five pathways representing pathways proliferation and oncogenetic drivers and hence tumor aggressiveness (*8*), see figure 4 below.

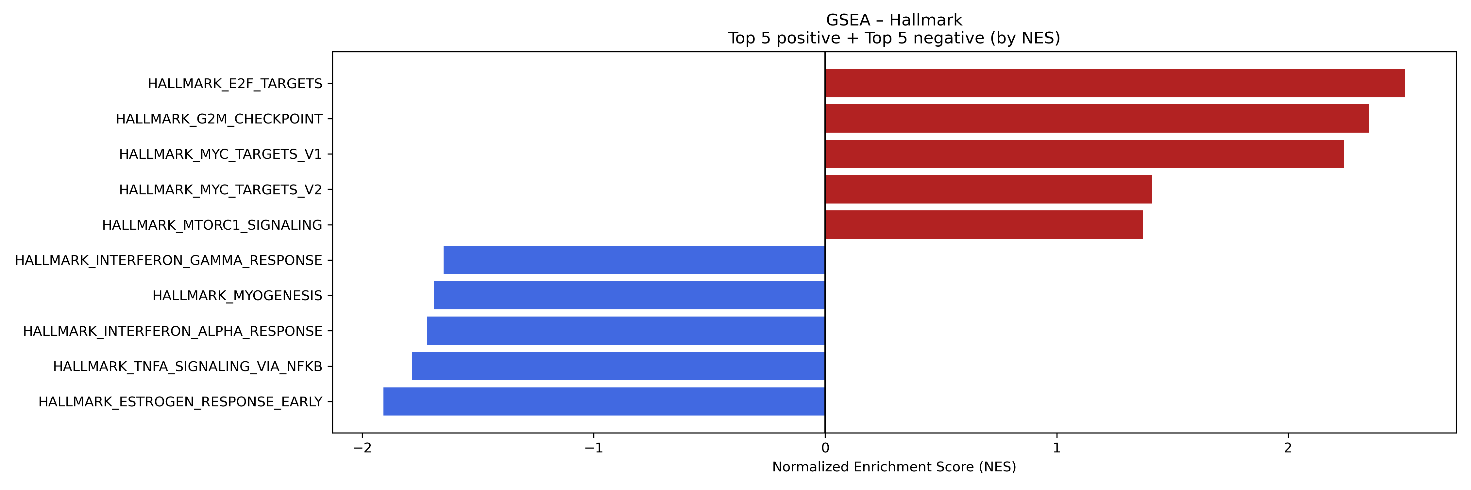

Supplementary Figure 3: Pathway enrichment analysis. Illustration of the top 5 ranked pathways between higher AmygAc and lower AmygAc. Blue indicating attenuated pathways and red increased expression in higher AmygAc patients.

**xCell Analysis**

xCell analysis revealed tendencies towards stress-associated differences in the enrichment of multiple immune cell-type gene signatures. These differences reflect relative changes in transcriptional programs associated with specific immune lineages rather than absolute differences in cell abundance. However, reduced enrichment score trends for CD8+ and CD4+ T cells, as well as Tregs subtypes can be observed. Due to high biological variance between samples, cell enrichment within samples of the groups is showing heterogeneity in the boxplots.

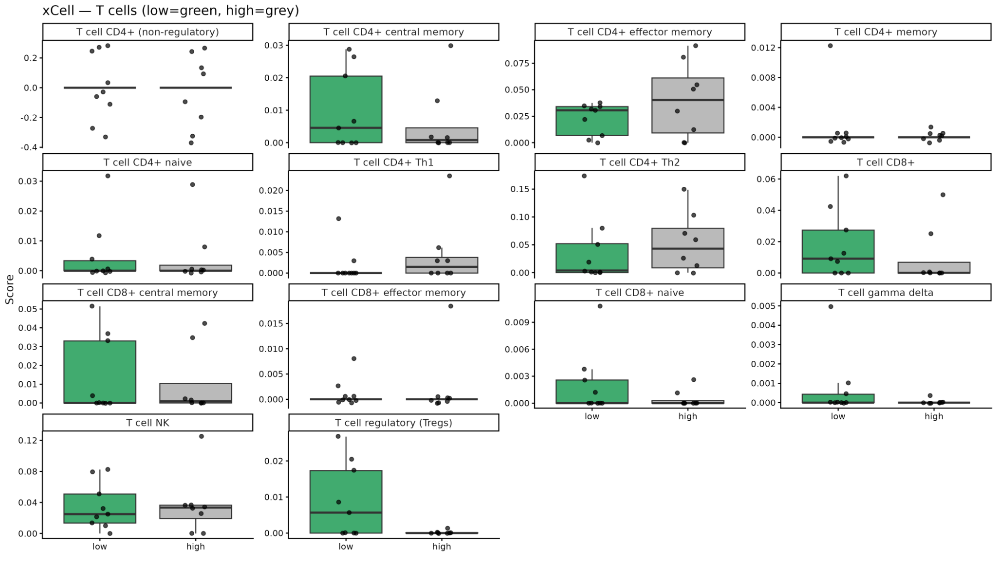

Figure 4: xCell enrichment scores for T-cell subsets in low- and high-stress samples. Boxplots display relative enrichment across samples (green: low stress; grey: high stress).

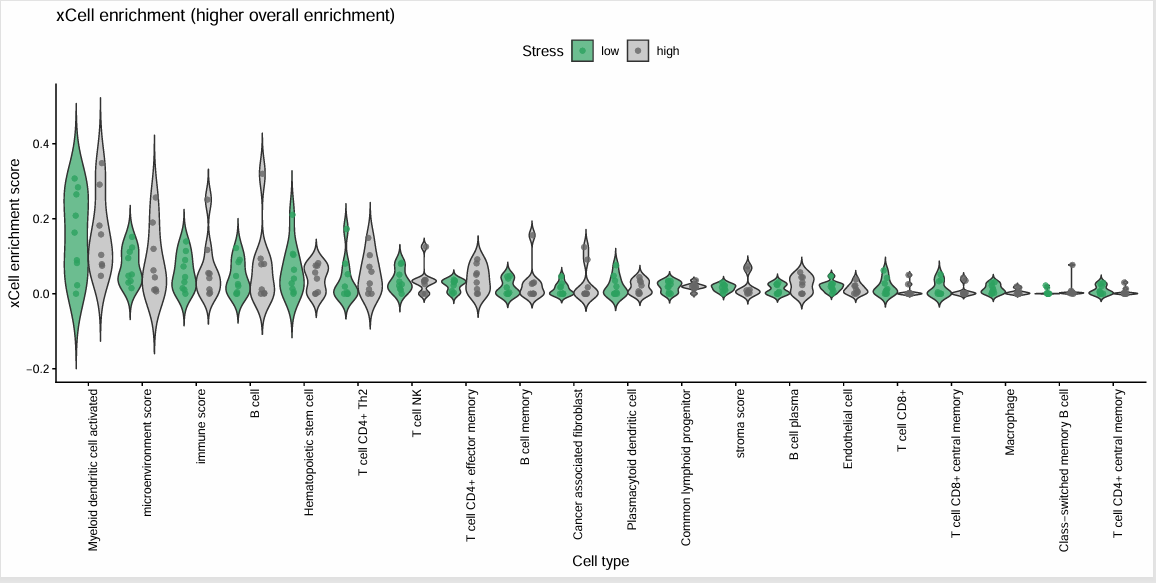

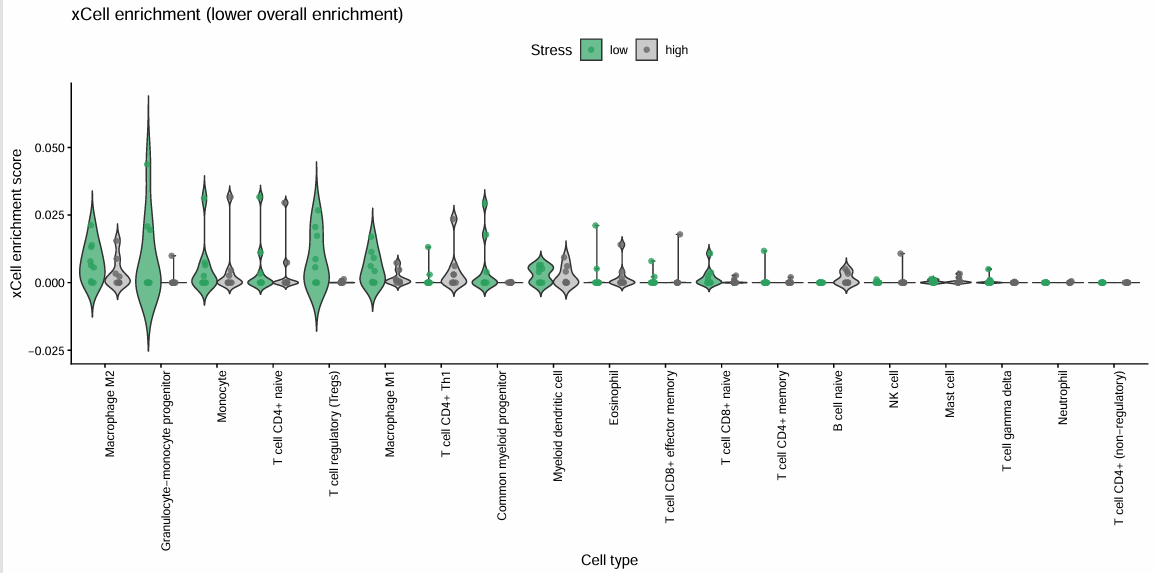

**MP-IF**

Figure 5: xCell enrichment scores for CellTypes provided by xCell in low- and high-stress samples. Violinplots display relative enrichment across samples (green: low stress; grey: high stress).

As we found differences between CTL density in total tissue (main manuscript) and TIME being significantly lower in the higher AmygAc group, we wanted to investigate the T-cell exhaustion marker PD-1 and PD-L1, expressed by tumor cells.

No differences were found between the stress groups for PD-L1 (P = 0.44) or PD-1 (P = 0.65) in total tissue or sub-regions (supplementary fig.2). The interactions between CTL and PD-L1 in the stressed groups are therefore not driven by a higher expression of exhausted CTL (measured by PD-1, and other exhaustion markers e.g. CTLA4 using transcriptomics analysis, Supp fig.1 and supp excel) and neither by amplified expression of PD-L1 by the tumor cells.

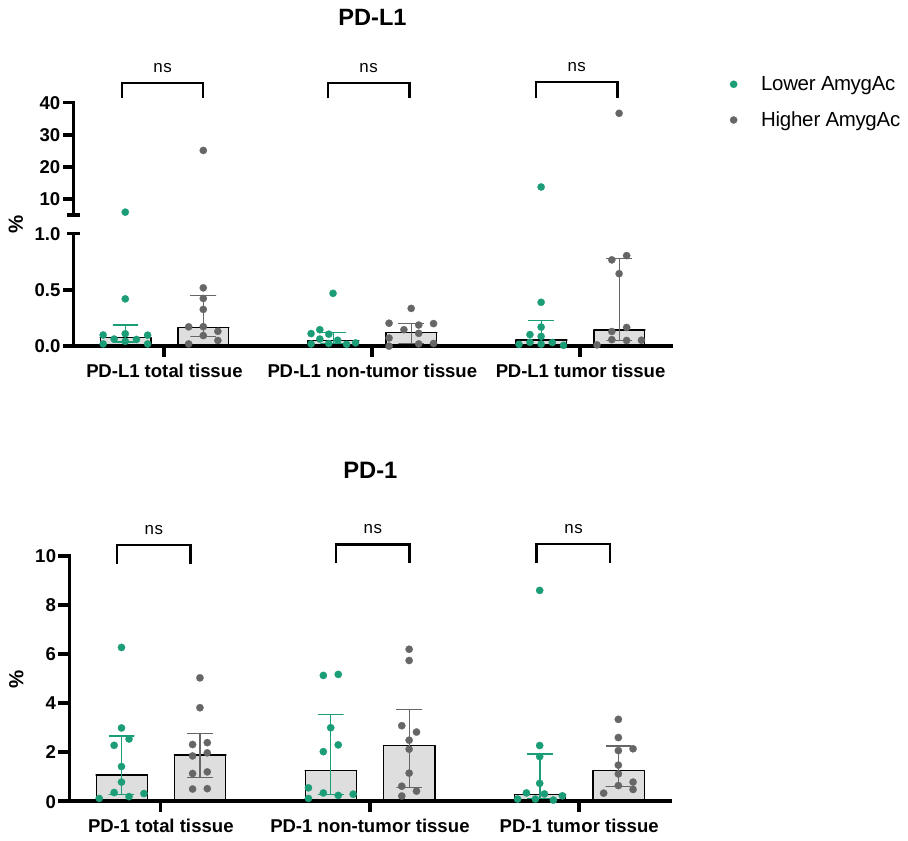

Supplementary Figure 6: Comparison of overall PD-L1 (above) and PD-1 (below) expressed as amount in percent between higher amygdala activity and lower amygdala activity group in total tissue (left), non-tumor tissue (center) and tumor tissue (right).

Also in the immunotherapy cohort, we neither observed differences in PD-L1 levels on tumor or immune cells nor differences in the combined PD-L1 Score between high and low AmygAc patients or respectively responders and non-responders (see supplementary table 2).

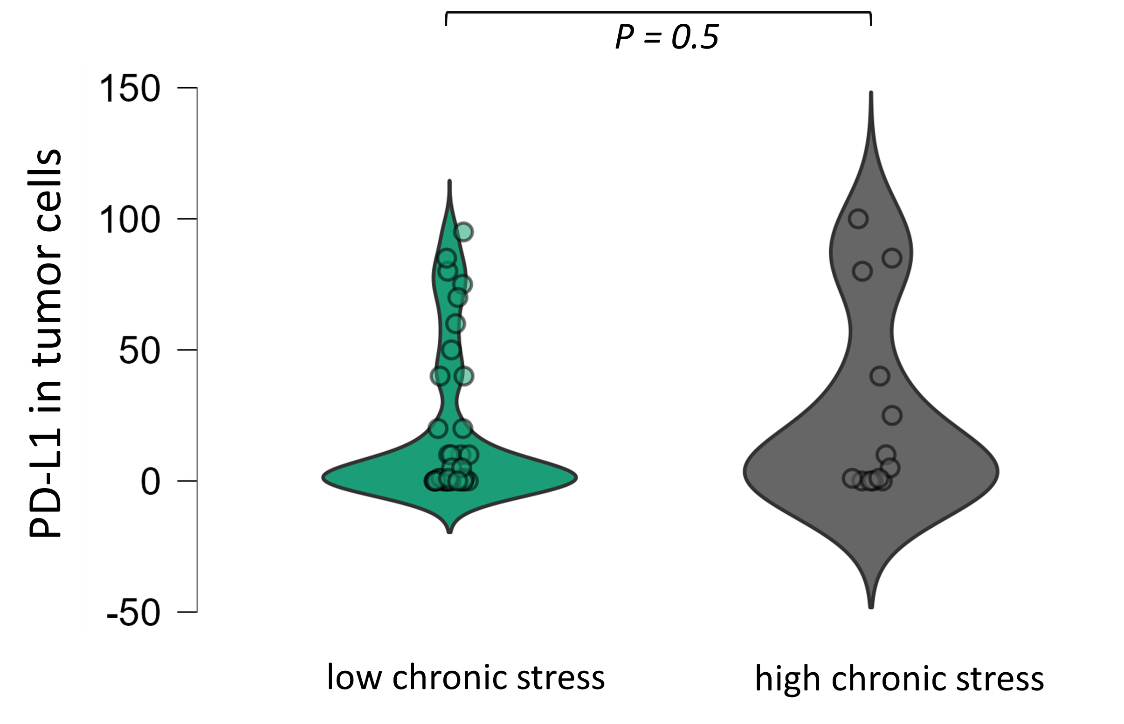

Supplementary Figure 7: Representative graph of the comparison of PD-L1 levels (%) on tumor cells in patients who underwent immunotherapy between higher amygdala activity and lower amygdala activity

PD-L1 levels in Immunotherapy Cohort

|  | **PD-L1 in tumor cells** | | **PD-L1 in immune cells** | | **Combined PD-L1 Score (CPS)** | |
| --- | --- | --- | --- | --- | --- | --- |
|  | **Median (IQR)** | **Mann-Whitney-U test P-value** | **Median (IQR)** | **Mann-Whitney-U test P-value** | **Median (IQR)** | **Mann-Whitney-U test P-value** |
| AmygAc high | **1.0 (0-20)** | **0.5** | **1.5 (0-8.75)** | **0.9** | **2.0 (0-25.75)** | **0.6** |
| AmygAc low | **5.0 (0-40)** |  | **1.0 (0-5.0)** |  | **12.5 (0-50)** |  |

**Supplementary Table 2:** Comparison of PD-L1 levels on tumor cells, immune cells and the combined PD-L1 Score in patients with high and low AmygAc treated with anti-PD-1/PD-L1 immunotherapy

Summarizing, we did not observe differences in PD-L1 or PD-1 levels between higher AmygAc and lower AmygAc neither in the therapy naïve nor in the immunotherapy treated cohort. Hence, cytotoxic T cells engage more frequently with PD-L1 in stressed individuals independently of total PD-L1 expression and coinciding with a lower number of CTLs total tumor tissue.
